# MiREDiBase: a manually curated database of validated and putative editing events in microRNAs

**DOI:** 10.1101/2020.09.04.283689

**Authors:** Gioacchino P. Marceca, Rosario Distefano, Luisa Tomasello, Alessandro Lagana, Francesco Russo, Federica Calore, Giulia Romano, Marina Bagnoli, Pierluigi Gasparini, Alfredo Ferro, Mario Acunzo, Qin Ma, Carlo M. Croce, Giovanni Nigita

**Affiliations:** Department of Clinical and Experimental Medicine, University of Catania, Catania, Italy; Department of Cancer Biology and Genetics and Comprehensive Cancer Center, The Ohio State University, Columbus, OH, USA; Department of Genetics and Genomic Sciences, Icahn School of Medicine at Mount Sinai, New York, NY, USA; Disease Systems Biology, Novo Nordisk Foundation Center for Protein Research, Faculty of Health and Medical Sciences, University of Copenhagen, Copenhagen, Denmark; Division of Pulmonary Diseases and Critical Care Medicine, Virginia Commonwealth University, Richmond, VA, USA; Fondazione IRCCS Istituto Nazionale dei Tumori (INT), Milan, Italy; Faculty of Health and Medicine, University of Newcastle, Newcastle, New South Wales, Australia; Department of Biomedical Informatics, College of Medicine, The Ohio State University, Columbus, OH, USA

## Abstract

MicroRNAs (miRNAs) are regulatory small non-coding RNAs that function as translational repressors. MiRNAs are involved in most cellular processes, and their expression and function are presided by several factors. Amongst, miRNA editing is an epitranscriptional modification that alters the original nucleotide sequence of selected miRNAs, possibly influencing their biogenesis and target-binding ability. A-to-I and C-to-U RNA editing are recognized as the canonical types, with the A-to-I type being the predominant one. Albeit some bioinformatics resources have been implemented to collect RNA editing data, it still lacks a comprehensive resource explicitly dedicated to miRNA editing. Here, we present MiREDiBase, a manually curated catalog of editing events in miRNAs. The current version includes 3,059 unique validated and putative editing sites from 626 pre-miRNAs in humans and three primates. Editing events in mature human miRNAs are supplied with miRNA-target predictions and enrichment analysis, while minimum free energy structures are inferred for edited pre-miRNAs. MiREDiBase represents a valuable tool for cell biology and biomedical research and will be continuously updated and expanded at https://ncrnaome.osumc.edu/miredibase.

## Introduction

MiRNAs are the most studied class of small non-coding RNAs involved in gene expression regulation. According to the canonical miRNA biogenesis pathway, miRNAs are initially transcribed into primary transcripts (pri-miRNAs) that present hairpin structures and undergo a double RNase III-mediated processing^1^. The first step occurs within the nucleus, where the Drosha-DGCR8 enzymatic complex cleaves pri-miRNAs into ~70 nucleotide long transcripts. These typically maintain the stem-loop conformation and represent the precursors of miRNAs (pre-miRNAs). Pre-miRNAs are then exported to the cytoplasm, where they are ultimately processed by Dicer into ~22 nucleotides long single-stranded RNAs (mature miRNAs)^1^. These can be found as −5p or −3p forms, depending on which miRNA’s arm they derive from^1^. To date, it is estimated that more than 1,900 pre-miRNAs are expressed in humans, giving rise to over 2,600 different mature miRNAs^2^.

MiRNAs are important modulators of gene expression^3,4^. The rule underlying their inhibitory activity over translation consists of a thermodynamically stable base pairing between a specific miRNA region, termed “seed region,” and a complementary nucleotide sequence of an mRNA, termed “miRNA responsive element” (MRE), causing the enzymatic degradation of the targeted transcript^3,4^. Conventionally, the seed region consists of nucleotides 2-8 located at the 5’ end of miRNAs and is usually assumed to interact with MREs included within the 3’ untranslated region (3’UTR) of target mRNAs^3,4^. MiRNAs take part in a vast range of physiological processes, including cell cycle control^5^, angiogenesis^6^, brain development^7^, behavioral changes, and cognitive processes^8^. Conversely, dysregulations of their expression or mutations in miRNA seed regions/MREs often lead to several pathologies, including tumors^5,9^.

RNA editing consists of the co- or post-transcriptional enzymatic modification of a primary RNA sequence through single-nucleotide substitutions, insertions, or deletions^10^. Recent transcriptome-wide analyses have revealed the pervasive presence of RNA editing in the human transcriptome. Currently, the adenosine-to-inosine (A-to-I) and cytosine-to-uracil (C-to-U) RNA editing are considered the canonical editing types^11^, with the A-to-I type being the most prevalent one^12–15^.

A-to-I RNA editing is catalyzed by enzymes of the Adenosine Deaminase Acting on RNA (ADAR) family, specifically ADAR1 (two isoforms) and ADAR2 (one predominant isoform)^16^. A-to-I RNA editing frequently occurs in non-coding transcripts, including pri-miRNA transcripts, and shows tissue-dependent patterns^12,15,17^. C-to-U RNA editing is catalyzed by enzymes of the Apolipoprotein B mRNA Editing Enzyme Catalytic Polypeptide (APOBEC) family, APOBEC1 and APOBEC3, at least in the context of mRNAs^18^. However, to date, no proof has been reported concerning the role of APOBECs in miRNA editing, and only a few studies have discussed this editing type in miRNAs^19,20^.

Editing of pri-miRNA exerts significant effects on miRNA biogenesis and function, with profound implications in pathophysiological processes, such as the progression of neurodegenerative diseases and cancers^16,21^. For instance, the editing of pri-miRNAs could induce a local structural change that prevents Drosha from recognizing the hairpin conformation, averting its cleavage, and allocating the edited pri-miRNAs to degradation^16^. Differently, editing events falling within the mature miRNA region not inducing the pre-miRNA suppression generate miRNAs diversified in their primary sequence, subsequently causing a change in miRNAs’ target repertoire (miRNA re-targeting)^16^.

Given the extensive number of high-confidence RNA editing sites retrieved so far and their relevance in the biomedical field, several efforts have been made to develop online resources capable of summarizing, contextualizing, and interpreting such data. These include databases like DARNED^22^, RADAR^23^, and REDIportal^24^, and more complex resources such as TCEA^25^. However, no dedicated online resources have been explicitly implemented for the study of miRNA editing until now. Here, we present MiREDiBase, the first comprehensive and integrative catalog of validated and putative miRNA editing events. MiREDiBase is manually curated and provides users with valuable information to study edited miRNAs as potential disease biomarkers.

## Results

### Data Collection

The MiREDiBase data processing workflow is depicted in Fig. 1. We first explored the PubMed literature by searching for specific keywords, such as “microRNA editing” and “miRNA editing,” narrowing the temporal range between 2000 and 2019. Retrieved articles were then manually filtered, discarding those not containing information on miRNA editing. Editing events detected or validated by targeted methods were included in the database and considered as authentic modifications. Among editing events detected through wide-transcriptome methods, we retained those established as “reliable” or “high-confidence” by the authors, classifying them as putative modifications. Statistical significance was taken into consideration when possible, eventually maintaining only significant editing events. We did not consider enzyme perturbation experiments as validation methods. For putative edited pre-miRNA sequences with no official miRNA name, e.g., “Antisense-hsa-mir-451” in Blow et al.^26^, we employed the BLASTN tool to generate alignments between the putative pre-miRNA sequence and miRBase’s pre-miRNA sequences (v22)^2,27^. Only perfect matches were retained and provided with their respective official name, as indicated by miRBase. In case editing positions were presented in the form of coordinates of previous genomic assemblies (i.e., hg19/GRCh37), these were converted to the hg38/GRCh38 assembly using the University of California Santa Cruz (UCSC) *liftOver* tool^28^. Editing sites associated with miRBase’s dead-entries were discarded.

**Figure 1.**
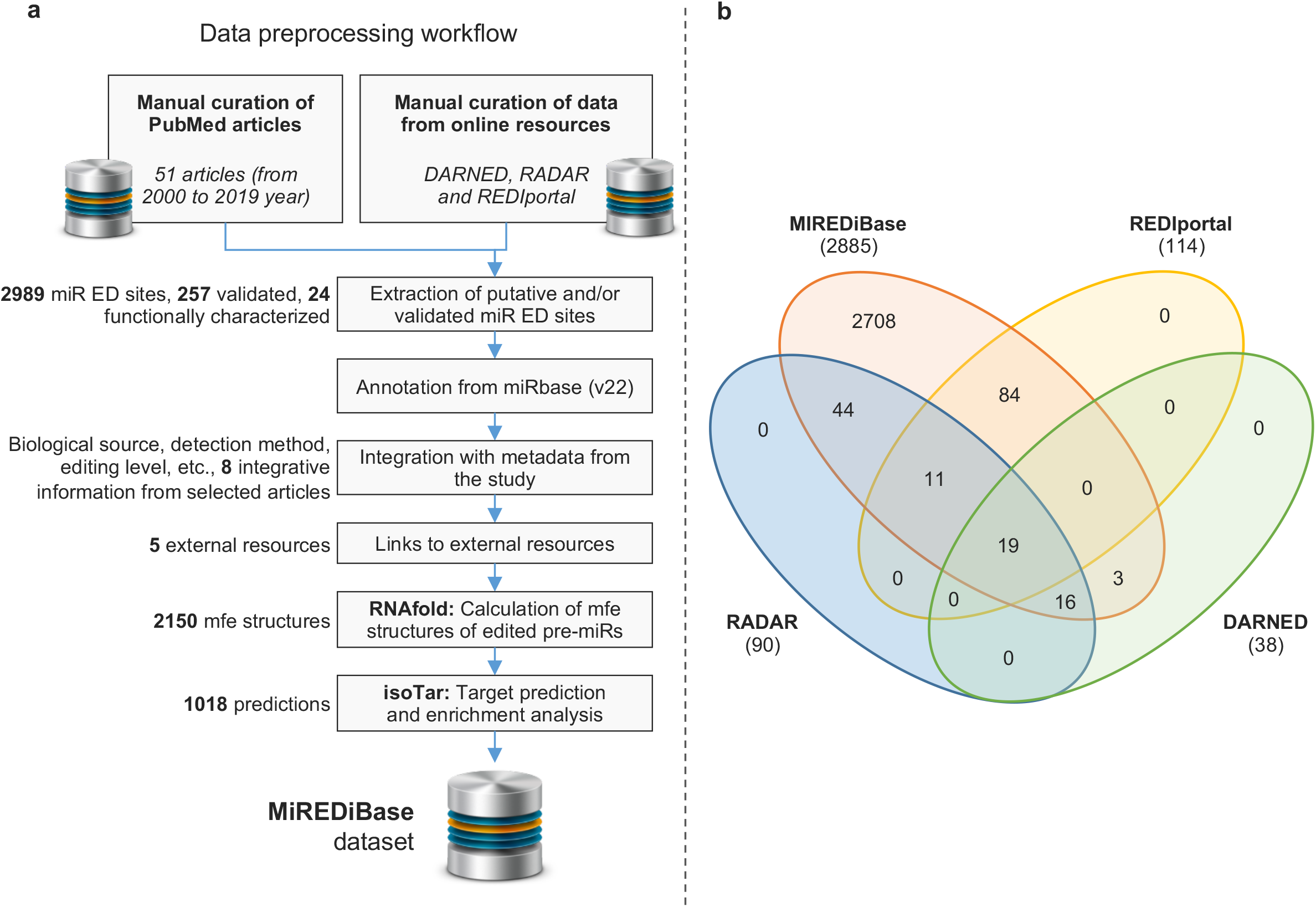
MiREDiBase’s data pre-processing workflow. (A) Chart is showing the workflow underlying miRTarBase. (B) Venn diagram showing the intersection of MiREDiBase with the three most prominent online repositories of A-to-I RNA editing events: DARNED, RADAR, and REDIportal. Data used for the intersection were exclusively relative to miRNA editing. The data were filtered for dead entries, opposite strands, and misassigned miRNAs prior to comparison.

In the second step, we expanded our search by employing the three most prominent online resources for A-to-I events available at present: DARNED^22^, RADAR^23^, and REDIportal^24^. Resources were manually screened, removing editing sites associated with dead entries and opposite strands. Editing sites falling into misassigned miRNAs in the hg19 genomic assembly (i.e., miRNAs of the hsa-mir-548 family and hsa-mir-3134 present in DARNED) were momentarily excluded from the database. The retained data were then integrated into the initial dataset.

### Database Content and Statistics

Considering the recent knowledge about genomic differences and similarities among primates, we retained data from *Homo sapiens* and three primate species (*Pan troglodytes*, *Gorilla gorilla*, and *Macaca mulatta*). In particular, the current version of MiREDiBase includes 2,989 validated and putative unique A-to-I (2,885) and C-to-U (104) editing events occurring in 571 human miRNA transcripts (Fig. 2a, 2b, Supplementary Data Set 1) and 70 unique A-to-I (46) and C-to-U (24) editing events taking place in 55 primate miRNA transcripts (Supplementary Figs. S1, S2a, S2b, Supplementary Data Set 1). Overall, 909 (29.7%) editing events occur outside of the pre-miRNA sequences, 971 (31.7%) within pre-miRNA sequences, outside of the mature sequence, and 1,179 (38.6%) within mature miRNA sequences (Fig. 3a, Supplementary Fig. S3a, Supplementary Data Set 1). These data were manually extracted from 51 original papers (Supplementary Table S1), which refer to 256 biological sources (Supplementary Tables S2-5).

**Figure 2.**
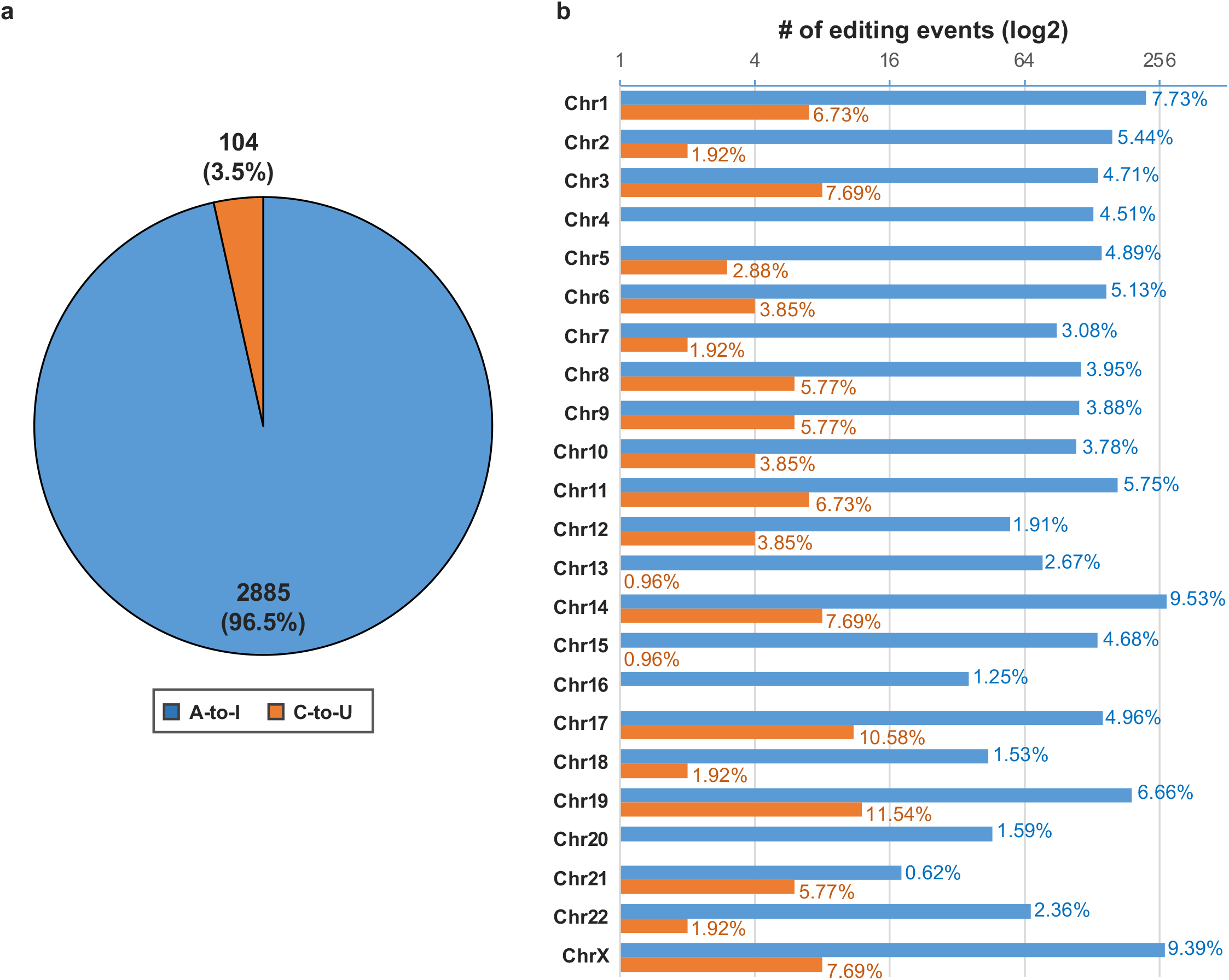
Descriptive statistics about human data in the current version of MiREDiBase. The number of unique A-to-I and C-to-U editing events reported from human tissues. (b) Distribution of A-to-I and C-to-U miRNA editing events per chromosome in the human genome. The percentage in each bar represents the percentage of the specific editing type (A-to-I or C-to-U) per chromosome, calculated respect the total number of that type editing event (A-to-I or C-to-U).

**Figure 3.**
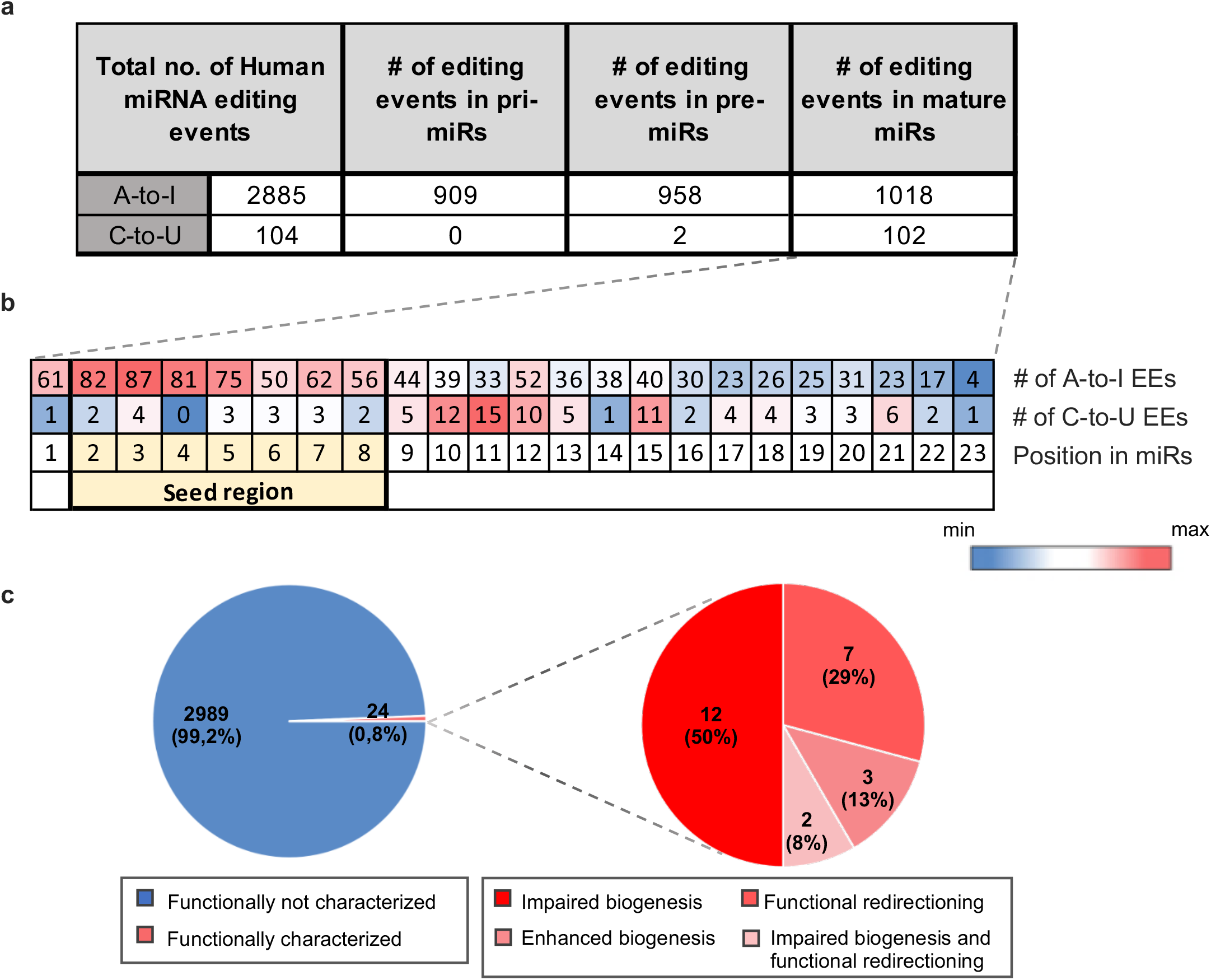
Editing events distribution in human microRNAs. (a) Distribution of A-to-I and C-to-U editing events across the three different regions of primary human miRNA transcripts. Heatmap of the distribution of A-to-I and C-to-U falling into human mature miRNAs across nucleotide positions. In each position are presents the number of editing events identified. The canonical seed position is highlighted in yellow. Editing sites falling in positions 24-27 were not shown in the mature sequence. EE=Editing Events. (c) Pie charts showing the fraction of functionally characterized editing events in human miRNAs.

Human editing sites in MiREDiBase are distributed across several genomic positions throughout the human genome, covering most chromosomes (Fig. 2b). However, of the 2,989 unique editing sites, only 257 (8.6%) have been validated by low-throughput methods or ADAR expression perturbation experiments. The majority of such events fall into clustered miRNAs located in chromosomes 14 (9.5% A-to-I; 7.7% C-to-U), chrX (9.4% A-to-I; 7.7% C-to-U), chr1 (7.7% A-to-I; 6.7% C-to-U), and chr19 (6.7% A-to-I; 11.5% C-to-U), respectively. Such a phenomenon very likely depends on local structural elements and motifs in these primary transcripts that function as editing inducers^29,30^ and would deserve more in-depth investigations. For the vast majority, the functionality of miRNA editing events has currently remained undetermined. So far, only 24 editing sites (0.8%) were functionally characterized by appropriate techniques (Fig. 3c). Among these, twelve were demonstrated to impair miRNA biogenesis; seven cause functional re-targeting; three cause impaired biogenesis and functional re-targeting; two cause enhancement of biogenesis.

Concerning primates, the majority of data refer to macaque (*Macaca mulatta*), for which our database reports 40 A-to-I and 24 C-to-U editing sites (Supplementary Figs. S1, S2, Supplementary Data Set 1). Here, 26 (65%) A-to-I editing sites are conserved between human and macaque, whereas only 8 (33%) C-to-U sites are conserved between these two species. This figure might suggest that A-to-I editing of miRNA transcripts is more conserved than the C-to-U type; however, it might also be due to the current low number of C-to-U instances reported for both human and primates. Only three editing sites are reported for both chimpanzee (*Pan troglodytes*) and gorilla (*Gorilla gorilla*), occurring in one pre-miRNA transcript for each species (Supplementary Data Set 1). None of the editing sites from primates have been validated yet.

When looking at editing sites falling within mature miRNA sequences, data from MiREDiBase let emerge two distinct patterns for A-to-I and C-to-U editing in humans (Fig. 3b). Examining the A-to-I type, most edited sites (325 out of 1018, 31.9%) are located at positions 2-5 of the seed region. Other hotspots for A-to-I editing seem to be represented by positions 1, 6-9, and 12, which account for 325 more edited sites. In the case of C-to-U miRNA editing, most, modification sites are located outside of the seed region. In particular, 48 out of 104 edited sites (46.2%) are located at positions 10-12 and 15, whereas only 17 edited sites (16.3%) fall within the seed region. A very similar pattern can be observed in macaque (Supplementary Fig. S3b). To help users interpret and contextualize data, miRNA editing events occurring within pre-miRNA or mature miRNA sequences were supplied with *in silico* predictions. We computed 2,150 MFE pre-miRNA predictive structures using editing sites internal to pre-miRNA sequences and 1,018 miRNA-targeting predictions and enrichment analyses. In both cases, users have the opportunity to compare the edited miRNAs with their relative wild-type versions. Biological sources in MiREDiBase can be grouped into three main categories (Table 1). The “normal condition” group (human and primates) accounts for 92 different healthy tissues/organs analyzed for miRNA editing. Among these, 85 were obtained from adult individuals and seven from pre-natal developmental stages (Supplementary Tables S2 and S5). The “adverse condition” group (human only) is broadly represented by tumors, with 60 distinct oncological conditions and 62 different sample subtypes. The neurological disorders include four pathological conditions and six sample subtypes. The inflammatory condition, cardiovascular disease, and genetic disorder are currently the less representative classes, with two pathological conditions and three sample subtypes for the former and one pathological condition and sample type for the latter two, respectively (Supplementary Table S3). The “cell line” group (human only) accounts for 78 commercial cell lines and ten primary human cells cultured *in vitro.* Of the 78 commercial cell lines, 71 are malignant, while the remaining represent non-malignant conditions. Among the ten primary human cells, only one refers to a malignant condition, while nine represent normal conditions (Supplementary Table S4).

**Table 1.** Number of normal and adverse conditions and cell lines present in MiREDiBase. The table shows the number and percentage of normal conditions, adverse conditions, and cell lines currently included in MiREDiBase for the four species Homo sapiens, Macaca mulatta, Pan troglodytes, and Gorilla gorilla.

|  | Species |  |  |  |
| --- | --- | --- | --- | --- |
| Condition | Human | Macaque | Chimpanzee | Gorilla |
| **Normal condition** | **69** | **23** | **1** | **1** |
| adult tissues | 64 (92,8%) | 21 (91.3%) | 1 (100%) | 1 (100%) |
| pre-natal tissues | 5 (7,2%) | 2 (8.7%) | NA | NA |
| **Adverse condition** | **68** | **0** | **0** | **0** |
| oncological diseases | 60 (88,2%) | NA | NA | NA |
| sample subtypes | 62 | NA | NA | NA |
| neurological diseases | 4 (5,9%) | NA | NA | NA |
| sample subtypes | 6 | NA | NA | NA |
| inflammatory diseases | 2 (2,9%) | NA | NA | NA |
| sample subtypes | 3 | NA | NA | NA |
| cardiological diseases | 1 (1,5%) | NA | NA | NA |
| sample subtypes | 1 | NA | NA | NA |
| genetic diseases | 1 (1,5%) | NA | NA | NA |
| sample subtypes | 1 | NA | NA | NA |
| **Cell lines** | **88** | **0** | **0** | **0** |
| non-malignant (commercial) | 7 (8,0%) | NA | NA | NA |
| non-malignant (ex vivo) | 9 (10,2%) | NA | NA | NA |
| malignant (commercial) | 71 (80,7%) | NA | NA | NA |
| malignant (ex vivo) | 1 (1,1%) | NA | NA | NA |

### User Interface and Data Accessibility

MiREDiBase provides users an intuitive and straightforward web interface to access data, requiring no bioinformatics skills to perform accurate searches across the database. Users can explore MiREDiBase by interacting with the Search (Fig. 4) or the Compare module. Each module starts with a modal box by which users can filter miRNA editing sites.

**Figure 4.**
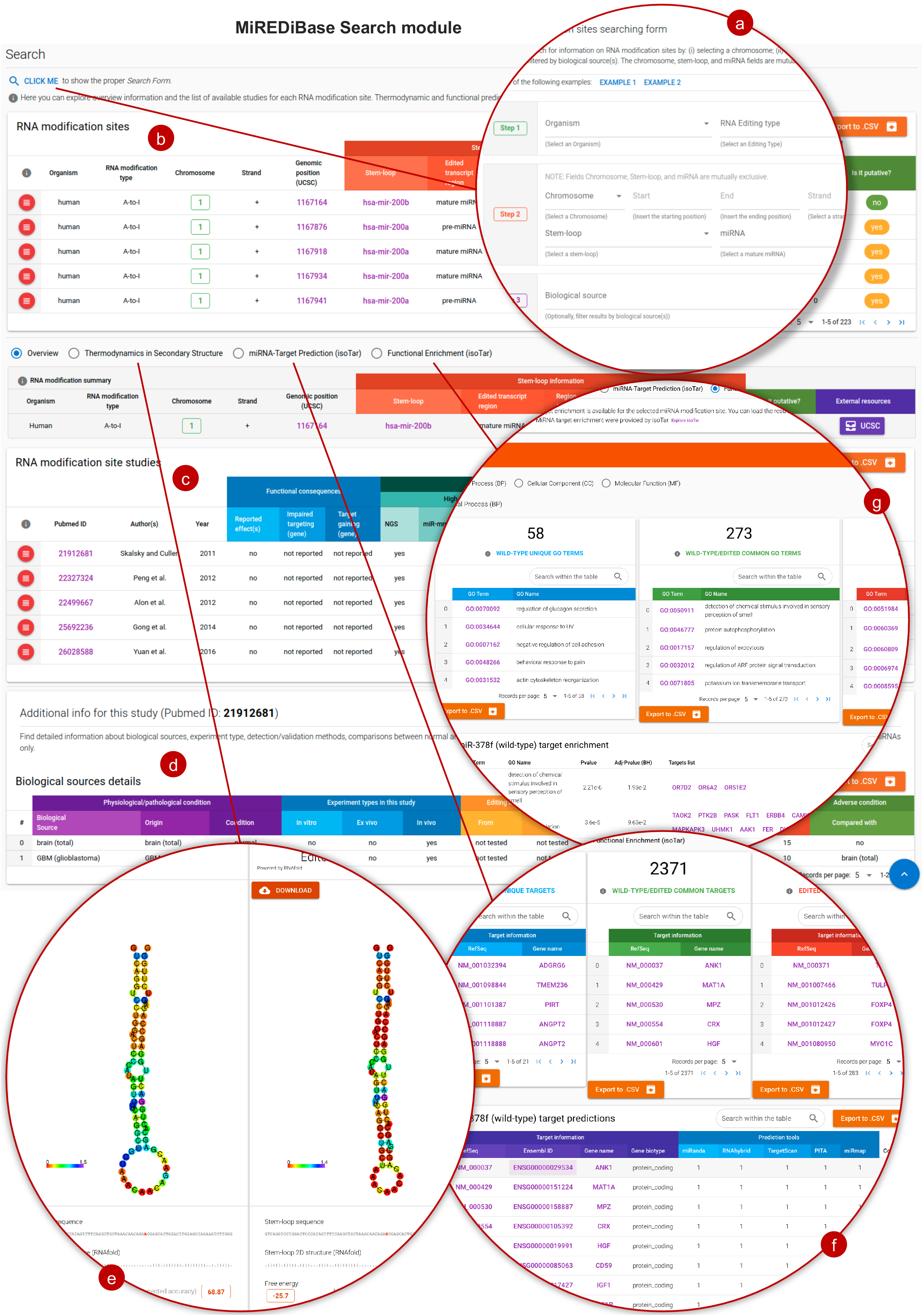
The MiREDiBase Search module. Users can filter out MiREDiBase data by exploiting the specific modal box (a). Then, they can dig into the data by interacting with the filtered editing sites (b). The editing site’s details (c) can be navigated by clicking on the red button placed on its left side. Additional resources include the list of biological sources in which the editing site has been identified (d), the thermodynamic comparison of the wild-type and edited pre-miRNA 2D structures (e), the miRNA-target predictions (f), and functional enrichment (G) data. Helpers and downloading buttons are provided throughout the module interface.

The Search module provides four filtering fields, including organism (e.g., Human), modification type (e.g., the A-to-I editing), genomic region (e.g., chromosome, pre-miRNA, or miRNA), and, optionally, biological source (e.g., BRCA – breast carcinoma). The “Search module” generates a table listing a set of editing sites supplied with essential information based on the selected filtering options. Reported information covers the organism, modification type, chromosome, strand, genomic position, pre-miRNA and mature miRNA relative positions, employed detection strategies, and whether the site is putative or validated. By clicking on the dedicated left-sided buttons, users can dig down to find supplementary information about each editing site. Here, the detection strategies information is expanded, categorizing the editing site as putative (i.e., only detected by high-throughput sequencing methods) or validated (i.e., authenticated by targeted methods), indicating the confidence level for each modification. Additional information covers publications, external resources, biological sources, 2D structures of edited and non-edited pre-miRNAs, miRNA-target predictions, and associated functional enrichment data (Fig. 4), which enable ready access to a putative biological interpretation. The results in each module can be easily downloaded through dedicated buttons. The Compare module aims at exploring differentially edited sites in adverse vs. normal conditions. It provides a set of essential information supplied with the editing level for each examined condition. Like the Search module, the Compare module allows users to filter out RNA editing sites by specifying the organism, modification type, disease, and pre-miRNA. All miRNAs reported in MiREDiBase are linked to their specific miRBase web page. Moreover, A-to-I genomic coordinates were mapped onto the UCSC hg38/GRCh38 genome assembly and available via the UCSC website. If applicable, editing sites provide links to external RNA editing resources, such as DARNED, RADAR, and REDIportal, to improve miRNA editing research.

To encourage users to familiarize themselves with our tool, MiREDiBase offers, throughout the website, helpers reporting explanations on how to interpret results, along with statistics and complete documentation on how to use each module. Advanced users can instead exploit the RESTful API, which provides a standalone web interface to explore available methods for extracting data, with the opportunity to embed RESTful API HTTP calls within users’ code (Fig. 5).

**Figure 5.**
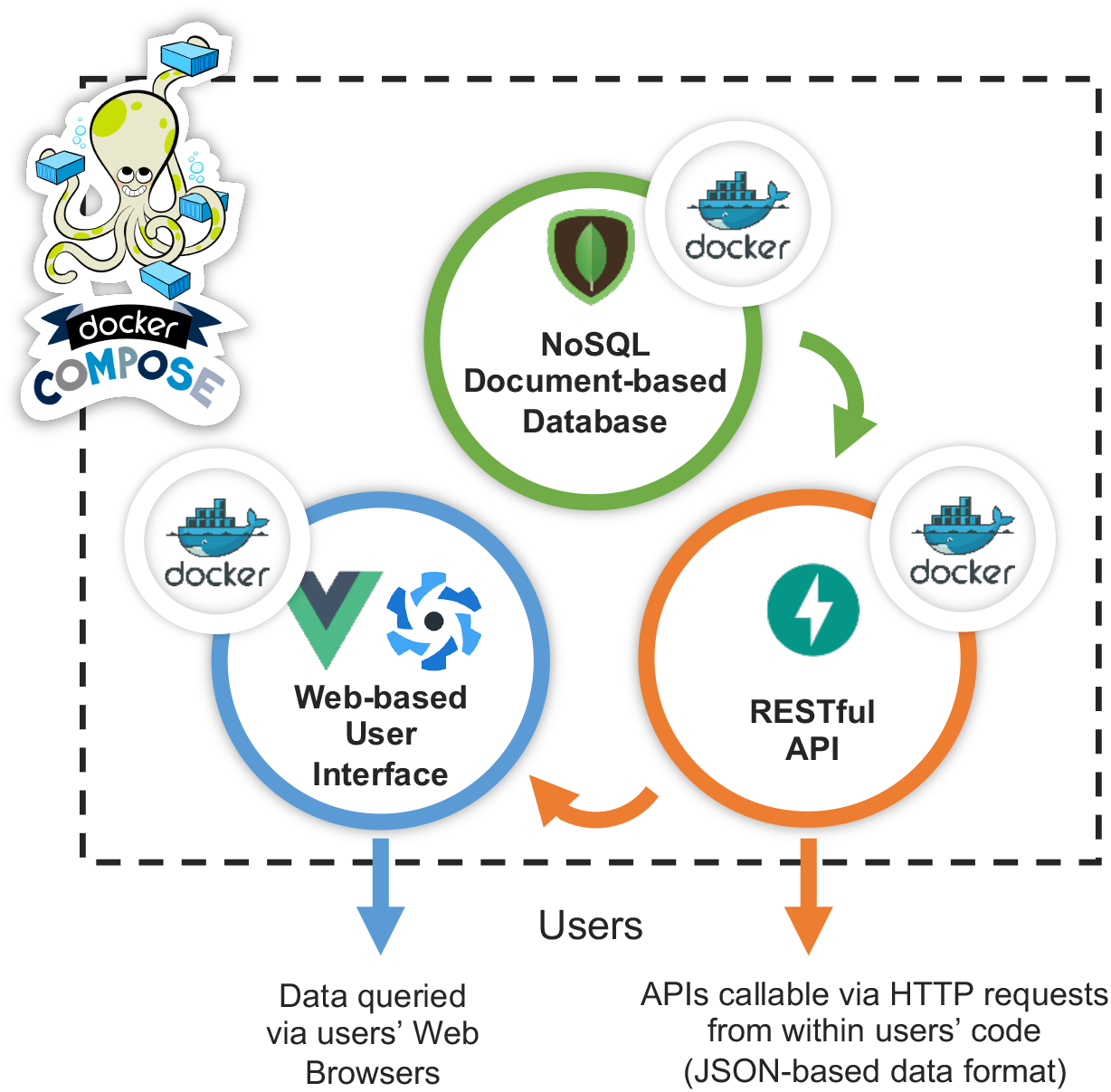
Overview of the MiREDiBase multi-containerized microservice architecture. The MiREDiBase platform provides users different ways to access its data: through a web browser (Web-based User Interface) and RESTful API (HTTP calls).

The MiREDiBase platform adopts a multi-containerized microservice architecture (Fig. 5), which provides user-friendly and efficient ways to access all manually collected data (see Methods section for more details).

## Discussion

At the beginning of the study on miRNA editing, Sanger sequencing represented the standard method to reliably identify editing events^31–33^. However, this low-throughput technique only enabled the detection of a relatively restricted set of editing sites. In later years, the employment of high-throughput sequencing (HTS) technologies and the design of *ad-hoc* bioinformatic pipelines have dramatically improved the computational identification of RNA editing events^34^, including those occurring in miRNAs.

Given the ever-increasing number of editing sites detected at a genome-wide scale, the need to create a comprehensive catalog of such modifications has become imperative. In light of this, Kiran and Baranov published DARNED, the first online repository providing centralized access to published data on RNA editing^22^. DARNED currently includes ~350,000 predicted RNA A-to-I editing sites from humans, mice (*Mus musculus*), flies (*Drosophila melanogaster*), and a few C-to-U instances. However, only a small portion of these modification events was manually annotated, and no information is provided about editing levels. DARNED’s last update dates back to 2012^35^.

In 2013, Ramaswami and Li presented RADAR, a rigorously annotated A-to-I RNA editing database containing manually curated editing sites^23^. Like DARNED, RADAR includes data from humans, mice, and flies and currently accounts for ~1.4 million editing sites, providing several useful information like tissue-specific editing level, conservation in other model organisms, and genomic context. RADAR does not include C-to-U editing data, and the update took place in 2014.

In 2017, Picardi and colleagues developed REDIportal, which today is the most extensive collection of RNA editing in humans, including more than 4.5 million A-to-I modification events detected across 55 body sites from thousands of RNA-seq experiments^24^. Moreover, with its last update, REDIportal also includes ∼90,000 putative A-to-I editing events from the mouse brain transcriptome and incorporates CLAIRE, a searchable catalog of RNA editing levels across cell lines^36^.

Although these three mentioned online resources are undoubtedly the most authoritative repositories of RNA editing events, none of them is strictly dedicated to miRNA editing. The vast majority of the editing events reported in these databases fall into mRNAs and long non-coding RNAs (lncRNAs), with only a minority occurring in miRNAs. Indeed, a few online resources have been lastly developed that partially focus on the effects of RNA editing on miRNA functionality. For instance, the Editome-Disease Knowledgebase (EDK)^37^ is a manually curated database that aims to link experimentally validated RNA editing events in non-coding RNAs to various diseases. However, this database currently contains only 16 validated A-to-I instances in miRNAs and does not provide any information about publications, position of editing sites, or detection/validation methods. The Cancer Editome Atlas (TCEA) is a powerful, user-friendly bioinformatics resource that characterizes more than 192 million editing events at ~4.6 million editing sites from approximately 11,000 samples across 33 cancer types recovered from The Cancer Genome Atlas^25^. However, TCEA is focused on editing events occurring in coding transcripts. From the miRNA standpoint, TCEA only allows users to predict A-to-I editing’s effects in the 3’ UTR of mRNAs in terms of miRNA-mRNA interactions. Analogous considerations apply for miR-EdiTar^38^, a database that exploits DARNED data to predict the potential effects of A-to-I editing over miRNA targeting.

To cover the gap between the fields of RNA editing and miRNA biology, we developed MiREDiBase, the first-of-its-kind database dedicated explicitly to miRNA modifications. In the current version, MiREDiBase includes more than three thousand A-to-I and C-to-U miRNA editing events manually collected from the literature, occurring in humans and primates. MiREDiBase allows users to consult the RNA secondary structure of both the wild-type and edited pre-miRNAs and infer the possible function of edited mature miRNA, based on the predicted targetome and subsequent functional analysis.

We implemented a user-friendly interface that allows users to track each search step to improve the user experience. Moreover, MiREDiBase includes a “Compare” section, which compares adverse versus normal conditions in a study-specific manner. Finally, the MiREDiBase platform relies on cutting-edge technologies, aiming at providing reliability and continuous operability. The platform represents an orchestration of different containerized services on top of Docker. Each service fulfills a specific purpose, such as a Web Application Service (Vue.js/Quasar - a Progressive JavaScript Framework), a RESTful API Service (FastAPI - a modern, high-performance, web framework for building secure APIs), and a Database Service (MongoDB - a NoSQL document-based database). The platform is designed to provide the smoothest and user-friendly experience to users.

We are aware that the lack of data on more commonly adopted model organisms and the inclusion of C-to-U RNA editing sites represent weaknesses in our work. The choice to include primates rather than other model species in this first release was motivated by the fact that primates present the highest genomic and transcriptomic similarity compared to humans^39^. Moreover, primates are recognized as excellent candidates to investigate epigenetic control of genome functions and are highly relevant for biomedical studies^39^. The choice to include putative C-to-U miRNA editing events was because this editing type is considered “canonical” among mammals. Indeed, previous Sanger-sequence validation of putative C-to-U editing sites in miRNAs found no evidence for real C-to-U miRNA editing^15,40^, letting hypothesize that such events were HTS artifacts. On the other hand, Negi et al. recently found and validated C-to-U editing at the fifth position of mature human miR-100, demonstrating that such an instance was functionally associated with CD4(+) T cell differentiation^20^. Given these controversies, we believe that collecting C-to-U miRNA sites with high consensus would serve to orientate future studies on this topic.

Besides expanding the database with published data, our main future goals are (i) to include editing events from other species, primarily model organisms like *Mus musculus* and *Drosophila melanogaster*, and (ii) adding other modification types. We believe that this will help interpret the functional roles of modified miRNA transcripts within the cell system. For example, after analyzing human brain samples for RNA editing events, Paul et al. unexpectedly found that a consistent percentage of miRNA editing events are non-canonical, especially C-to-A and G-to-U^11^. Similar data were reported by Wang and co-workers^41^, raising questions on whether these editing events exert essential function in neurons and if specific enzymes can catalyze such modifications. Likewise, miRNA methylation has recently caught the scientific community’s attention, being demonstrated to affect miRNA biogenesis^42^. However, the study of this phenomenon and its potential functional implications have remained widely unexplored. With continuous updating, we believe that MiREDiBase will gradually become a precious resource for researchers in the field of epitranscriptomics, leading to a better understanding of miRNA modification phenomena and their functional consequences.

## Methods

### Data Processing

Each editing event was supplied with essential information recovered from miRBase (v22), including the relative position within pre-miRNA and mature miRNA, genomic position, and pre-miRNA region (5’- or 3’-arm, or loop region). For editing events occurring outside the pre-miRNA sequence, we adopted the notation “pri-miRNA.” Editing events were then enriched with metadata manually collected from selected publications. Overall, we extracted eight different information types: detection/validation method, experiment type, biological source, correspondent condition (adverse or normal), comparison (pathological vs. physiological condition), editing level, enzyme affinity, and functional characterization.

The “detection method” does not specify the method adopted by authors to identify miRNA editing events. Instead, it indicates which kind of methodological approach (targeted, wide-transcriptome, or both) the authors selected for editing detection. Only in two cases, the method has been specified to highlight particularly sensitive and innovative approaches, i.e., miR-mmPCR-seq^15,43^ and RIP-seq^44^.

The “validation method” refers to methods confirming sequencing data, especially those obtained by wide-transcription approaches, including enzyme knock-down (only ADAR in the current version), knock-out, differential expression, and modification-specific enzymatic cleavage.

The “experiment type” specifies whether, in a particular study, individual editing events were identified *in vitro*, *in vivo*, or *ex vivo*. Editing events obtained by analyzing small RNA-seq data from The Cancer Genome Atlas (TCGA)^45^ or Genotype-Tissue Expression (GTEx) atlas^46^ were considered as detected in vivo. Editing events obtained by analyzing sequence libraries from the Sequence Read Archive (SRA) database^47^ were considered as detected *in vitro*, *in vivo*, or *ex vivo* depending on the library derivation.

The “pathological condition” specifies whether a miRNA editing event was detected in one or multiple diseases. For a given study, physiological and pathological conditions were compared to whether editing levels for an individual miRNA were simultaneously available for both conditions.

In studies with multiple editing level values per miRNA editing site, we considered only the minimum and maximum values, rounding them up by multiples of five (e.g., editing levels of 21.1% and 44% were rounded up to 20% and 45%, respectively). Whether a single value was reported for an individual miRNA, this was rounded, creating an interval of 5% (e.g., if a study reported the editing level as 13% for a specific editing site, the editing level was presented as “from 10% to 15%”).

Information concerning enzyme affinity (only ADARs in the current version) was retrieved whether authors carried out enzyme-transfection experiments causing enzyme overexpression. Finally, we annotated all the functionally characterized editing events with information regarding their specific biological function. In the event of functional re-targeting, validation methods were reported along with the set of validated lost and gained targets.

### Secondary Structure Prediction Analysis

We generated the minimum free energy (MFE) structures for all those pre-miRNAs subjected to editing and their wild-type (WT) counterparts. The double-stranded RNA structures were created by employing the *RNAfold* tool from the ViennaRNA package^48^ with default settings. Finally, we considered all editing sites occurring within the mature miRNA region to infer possible miRNA target re-direction as well as diversified biological functions.

### MiRNA-Target Prediction and Functional Enrichment Analyses

The miRNA-target prediction analysis, for both edited and WT miRNA, was achieved by using our web-based containerized application *isoTar*^49^, designed to simplify and perform miRNA consensus target prediction and functional enrichment analyses. For miRNA target predictions, we established a minimum consensus of 3. An adjusted P-value <0.05 was considered as a threshold for the functional enrichment analysis.

### Platform Design and Implementation

To achieve reliability and continuous delivery (short-cycle updates), we developed each lightweight, standalone, microservice on top of Docker (v19.03.12) (https://docs.docker.com)^50^. The platform itself consists of three microservices, which are orchestrated by Docker Compose (v1.26.0) (https://docs.docker.com/compose)^51^, a tool for managing multi-container-based applications. Each microservice provides a specific functionality: a web-based user interface (UI), a RESTful API for data retrieving, and a NoSQL document-based database for data storing. To offer users an engaging and responsive experience, we developed a high-performance platform which relies on cutting-edge open-source technologies, such as Quasar (v1.12.8) (https://quasar.dev)^52^ for the UI, FastAPI (v0.55.1) (a Python (v3.8.1) framework, https://fastapi.tiangolo.com)^53^ for the RESTful API, and MongoDB (v4.2.8) (https://www.mongodb.com)^54^ for the data storage. We have tested MiREDiBase on the following browser: Firefox (80+), Google Chrome (85+), Edge (85+), Safari (13+), and Opera (70+). MiREDiBase is freely accessible to the scientific community through the link: https://ncrnaome.osumc.edu/miredibase, without requiring registration or login.

## Data Availability

MiREDiBase is freely available at https://ncrnaome.osumc.edu/miredibase.

## Code Availability

The whole platform is openly available at https://github.com/ncRNAome-OSU/miredibase.

## Acknowledgements

We thank the Cancer IT Operation Group of The Ohio State University (OSU) and Thomas Moore for his valuable technical assistance. At the same time, we want to special thank Paolo Fadda from The Genomics Shared Resource (GSR) of OSU, together with Jing Jiang and Cankun Wang from Dr. Ma’s Laboratory (Department of Biomedical Informatics at OSU) for their helpful suggestions during the beta testing phase of MiREDiBase.

This work was supported by National Cancer Institute (National Institute of Helth) grant R35CA197706 to C.M.C. F.R. is supported by the BRIDGE - Translational Excellence Programme (bridge.ku.dk), Faculty of Health and Medical Sciences, University of Copenhagen, funded by the Novo Nordisk Foundation with grant agreement no. NNF18SA0034956 and grant agreement no. NNF14CC0001.

## Author contributions

C.M.C. and G.N. conceived, designed, and supervised the project; G.P.M., R.D., L.T., and G.N. designed the database; G.P.M. collected the data; R.D. designed and developed the platform; L.T., A.L., F.R., F.C., G.R., M.B., P.G., A.F., M.A., and Q.M. contributed to the beta-testing phase of the database. G.P.M., R.D., and G.N. wrote the first draft of the manuscript. L.T., A.L., F.R., F.C., G.R., M.B., P.G., A.F., M.A., Q.M., and C.M.C. participated in manuscript revision. All authors read and approved the final manuscript.

